# OrthoFinder: phylogenetic orthology inference for comparative genomics

**DOI:** 10.1101/466201

**Authors:** David M. Emms, Steven Kelly

## Abstract

Here, we present a major advance of the OrthoFinder method. This extends OrthoFinder’s high accuracy orthogroup inference to provide phylogenetic inference of orthologs, rooted genes trees, gene duplication events, the rooted species tree, and comparative genomic statistics. Each output is benchmarked on appropriate real or simulated datasets and, where comparable methods exist, OrthoFinder is equivalent to or outperforms these methods. Furthermore, OrthoFinder is the most accurate ortholog inference method on the Quest for Orthologs benchmark test. Finally, OrthoFinder’s comprehensive phylogenetic analysis is achieved with equivalent speed and scalability to the fastest, score-based heuristic methods. OrthoFinder is available at https://github.com/davidemms/OrthoFinder.

## Background

Determining the phylogenetic relationships between gene sequences is fundamental to comparative biological research. It provides the framework for understanding the evolution and diversity of life on Earth and enables the extrapolation of biological knowledge between organisms. Given the central importance of this process to multiple areas of biological research, a diverse array of methods have been developed that attempt identify these relationships given sets of user-supplied gene sequences [1–3]. The majority of these methods try to deduce phylogenetic relationships between gene sequences through heuristic analyses of pairwise sequence similarity scores (or expectation values) obtained from an all-vs-all BLAST [4] search, or accelerated alternatives to BLAST such as DIAMOND [5] or MMseqs2 [6]. Widely used methods include InParanoid [7], OrthoMCL [8], OMA [9] and OrthoFinder [10] all of which take different approaches to interrogating sequence similarity scores, and all of which produce different outputs—some identify orthogroups, some identify orthologs and paralogs, some do both. As they each adopt different approaches to analysing sequence similarity scores, each of the methods exhibit different performance characteristics on commonly used benchmark databases [1, 11].

Heuristic analysis of pairwise sequence similarity scores has historically been used to estimate the phylogenetic relationship between genes as it is readily computationally tractable. The central premise underlying their use is that higher scoring sequence pairs are likely to have diverged more recently than lower scoring sequence pairs. Thus, heuristic analysis of sets of pairwise sequence similarity scores can be used to estimate the phylogenetic relationships between sets of genes [7–9, 12, 13]. However, such score-based estimates of the phylogenetic relationship between genes are confounded by multiple factors. For example, variable sequence evolution rates between genes frequently leads to both false positive and false negative errors [14, 15] (Fig. 1). Such errors can be mitigated by analysis of phylogenetic trees of genes [16], as phylogenetic trees are able to distinguish variable sequence evolution rates (branch lengths) from the order in which sequences diverged (tree topology) (Fig. 1). Although an automated method for phylogenetic ortholog inference does not exist, curated databases of orthologs obtained from analysis of phylogenetic trees have been constructed [17–20]. These curated databases generally have higher ortholog detection accuracy than score-based approximate methods [1], but are limited to the species previously chosen for analysis by the database provider. Thus, there is a need for an automated method that has the enhanced accuracy of a phylogenetic method, but with the ease of use, speed and scalability of a score-based heuristic method.

**Figure 1.**
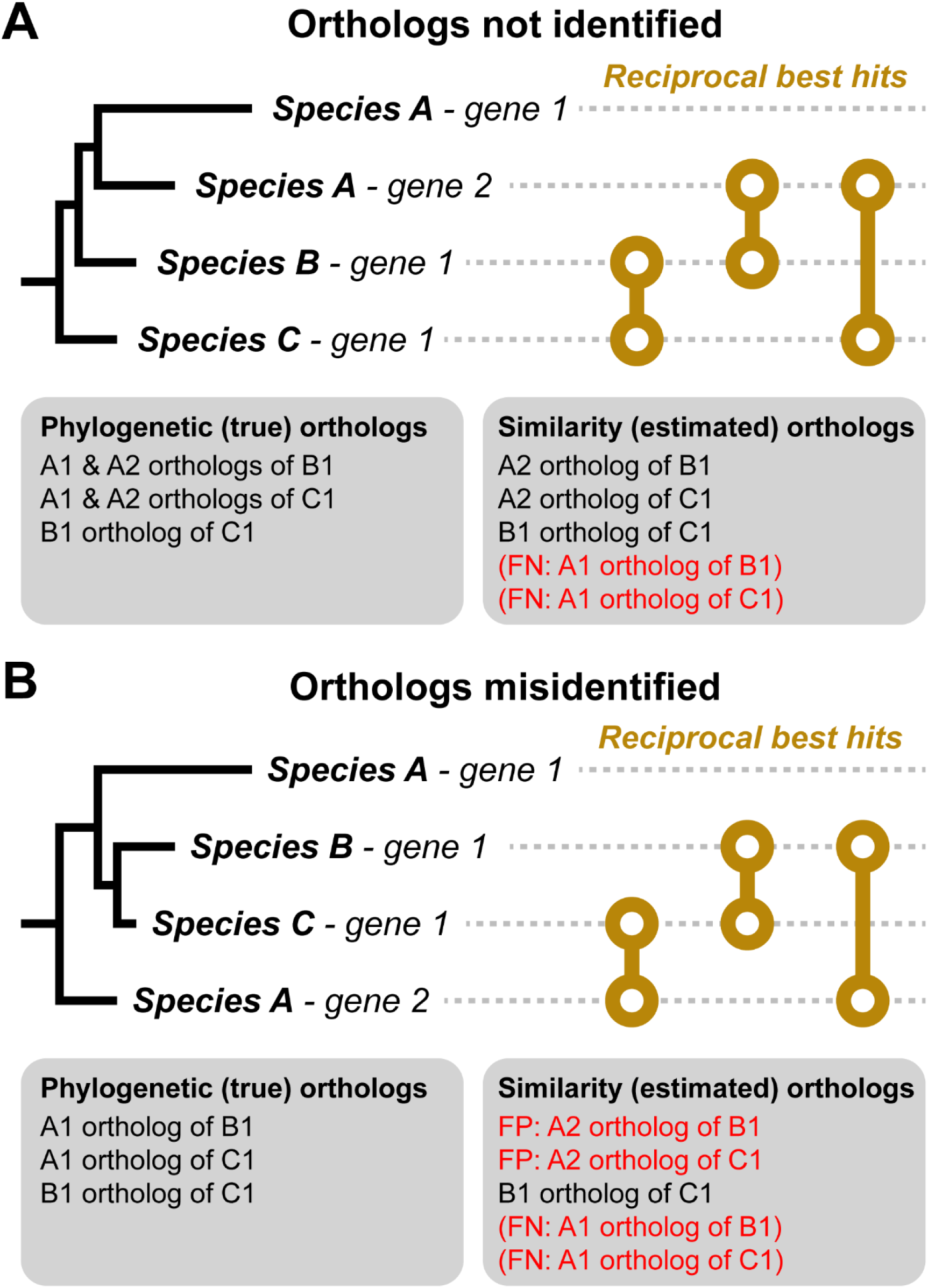
Pairwise similarity-score based ortholog inference can be misled by variable sequence evolution rates. A) Phylogenetic tree of a typical gene family. The correct orthologs of Species A – gene 1 are not identified. B) Species A – gene 2 is misidentified as the ortholog of the genes from Species B & C. Left-hand side: gene trees with branches to scale and the true orthology relationships, which can be determined from the gene tree. Right-hand side: Reciprocal best hits (RBH) based on gene similarity scores that are monotonic with branch length and the orthology relationships inferred from these scores using standard heuristics (orthologs inferred using RBHs and co-orthology identified from within species hits better than closest RBH [8, 45]). FP: false positive, FN: false negative.

While an automated method for phylogenetic orthology inference from gene sequences is an important goal, the implementation of such a method presents several technical challenges. These comprise:1) Inferring a complete set of gene trees for all genes of a given set of species in a time-scale that is competitive with score-based heuristic methods. 2) Automatically rooting these gene trees so that they can be correctly interpreted [21] without requiring the user to know the rooted species tree in advance. 3) Interpreting the gene trees to identify gene duplication events, orthologs and paralogs while being robust to processes such as gene duplication, loss, incomplete lineage sorting and gene tree inaccuracies. If these challenges could be addressed in a resource and time efficient manner, then such a phylogenetic method would provide a step change for orthology inference, enabling the transition from similarity-score based estimates of phylogenetic relationships to phylogenetically delineated phylogenetic relationships between genes.

Although a complete method for phylogenetic orthology inference from gene sequences does not exist, some of the challenges listed above have already been addressed in isolation by a range of bioinformatic methods. For example, there are a range of methods for identifying orthogroups of genes from user supplied gene sequences [8–10, 12, 22] and a wide variety of gene tree inference methods that can infer trees from these orthogroups [23–27]. Similarly, there is a range of methods for inferring orthologs from gene trees that also vary in terms of scalability and accuracy [28–31]. However, prior to the development of the version of OrthoFinder described in this paper, other critical challenges had no existing solutions. For example, there was no automated method to infer a set of rooted gene trees given only gene sequences as input. Equally, methods to infer orthologs from gene trees did not exist that were robust to processes such as incomplete lineage sorting and gene tree inference error while also being scalable to the large scale analysis required for whole-genome orthology inference across hundreds of species. Thus, substantial technical challenges needed to be addressed to enable fully automated and efficient phylogenetic delineation of the phylogenetic relationships between genes.

Here, we present a major update to OrthoFinder that significantly extends the scope of the original method. The updated version of OrthoFinder identifies orthogroups as in the original implementation [10], but then uses these orthogroups to infer gene trees for all orthogroups and analyses these gene trees to identify the rooted species tree. The method subsequently identifies all gene duplication events in the complete set of gene trees and analyses this information in the context of the species tree to provide both gene tree and species tree level analysis of gene duplication events. Finally, the method analyses all of this phylogenetic information to identify the complete set of orthologs between all species and provide extensive comparative genomics statistics. The complete OrthoFinder phylogenetic orthology inference method is accurate, fast, scalable and customisable and is performed with a single command using only protein sequences as input.

## Results

### OrthoFinder algorithm overview and summary of results files

The OrthoFinder algorithm is described in detail in the Methods. In brief, it address the challenges identified above in five major steps: a) orthogroup inference; b) inference of gene trees for each orthogroup; c & d) analysis of these gene trees to infer the rooted species tree; e) rooting of the gene trees using the rooted species tree; and f-h) duplication-loss-coalescence (DLC) analysis of the rooted gene trees to identify orthologs and gene duplication events (mapped to their locations in both the species and gene trees) (Fig. 2). Thus starting from just gene sequences OrthoFinder infers orthogroups, orthologs, the complete set of gene trees for all orthogroups, the rooted species tree, all gene duplication events and computes comparative genomic statistics. To illustrate the standard outputs provided by an OrthoFinder analysis, a graphical example of the complete set of results produced by OrthoFinder for 10 metazoan species are shown in Figure 3A-H.

**Figure 2.**
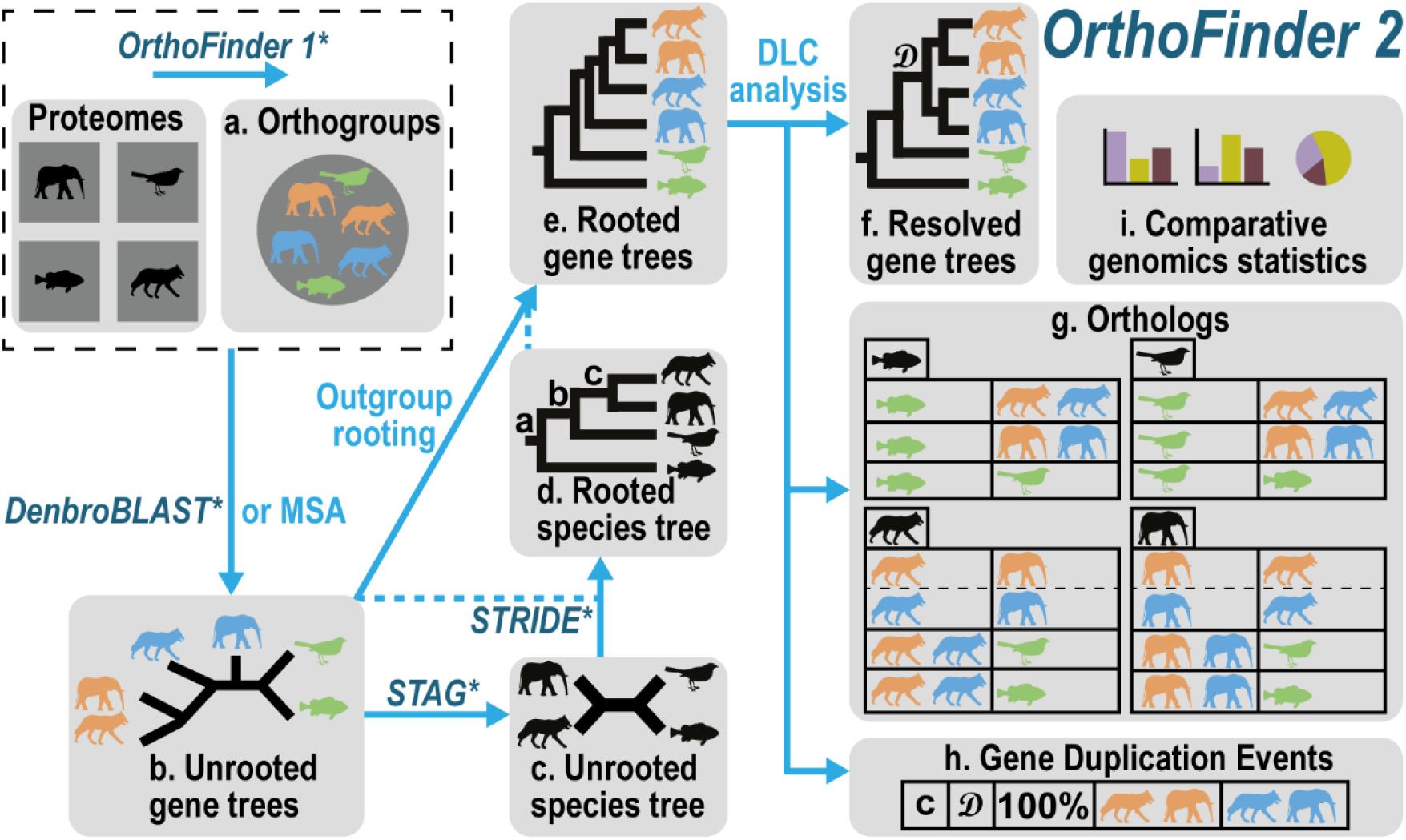
The OrthoFinder workflow: The method used for each step is shown by the arrow. Published algorithms are shown in italics and are followed by an asterisks. A dotted blue line connecting with a solid arrow indicates additional data that are used in order to carry out the transformation indicated by the solid arrow. MSA = multiple sequences alignment based tree inference, DLC = Duplication-Loss-Coalescence. The results tables in (g) illustrate the orthologs for the genes in each input species (four main boxes). The horizontal divisions within these show the orthologs for each individual species-pair. The gene duplication table in (h) shows the location of the duplication mapped to the species tree, the location in the gene tree, the percent retention of the duplicate genes in the samples species and the genes descended from the gene duplication event.

**Figure 3.**
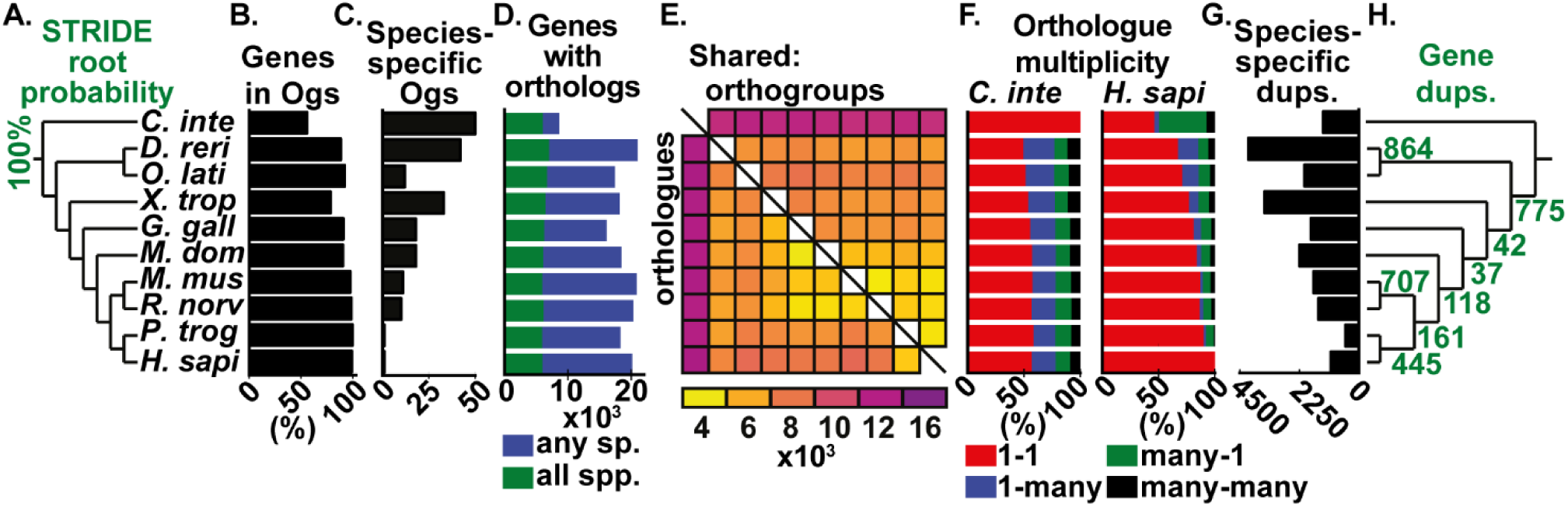
Summary of OrthoFinder analysis of a set of Chordata species: *Ciona intestinalis, Danio rerio, Oryzias latipes, Xenopus tropicalis, Gallus gallus, Monodelphis domestica, Mus musculus, Rattus norvegicus, Pan troglodytes* & *Homo sapiens*. Bar charts and heat map contain data for each species, aligned to the corresponding species in the tree in (A). A) The species tree inferred by STAG and rooted by STRIDE B) Percentage of genes from each species assigned to orthogroups. C) The number of species-specific orthogroups D) Number of genes with orthologues in any/all species. E) Heat map of the number of orthogroups containing each species-pair (top right) and orthologues between each species (bottom left) F) Orthologue multiplicities for two species, *C. intestinalis* and *H. sapiens*, with respect to all other species G) Number of gene duplications events on each terminal branch of the species tree. H) Number of duplications on each branch of the species tree and retained in all descendant species. Abbreviations: OG=orthogroup, sp.=species, spp.=species (plural), dups.=gene duplication events

The default, and fastest, version of OrthoFinder uses DIAMOND [23] for sequence similarity searches. These sequence similarity scores provide both the raw data for orthogroup inference [10] and for gene tree inference of these orthogroups using DendroBLAST [23]. The default implementation of OrthoFinder been designed to enable a complete analysis with maximum speed and scalability using only gene sequences as input. However, OrthoFinder has also been designed to allow the use of alternative methods for tree inference and sequence search to accommodate user preferences. For example, BLAST [4] can be used for sequence similarity searches in place of DIAMOND. Similarly, gene trees do not need to be inferred using DendroBLAST. Instead, OrthoFinder can automatically infer multiple sequence alignments and phylogenetic trees using any user preferred multiple sequence alignment and tree inference method. Moreover, if the species tree is known prior to the analysis, this can also be provided as input, rather than inferred by OrthoFinder. Thus, while OrthoFinder is designed to require minimal inputs and computation, it can be tailored to suit the computational and data resources available to the user.

### OrthoFinder has the highest orthology inference accuracy

The accuracy of key component algorithms of OrthoFinder have been independently assessed both above and in dedicated publications [5, 10, 21, 23, 32]. To demonstrate the accuracy of the overall method, the orthologs identified by OrthoFinder using its default options, along with multiple different configurations, were submitted to the community supported *Quest for Orthologs* benchmarking server [1] (see methods for details of tests). The results of all of these analyses are shown in Fig. 4A-L, and supported by additional analyses in Supplementary Fig. 1 and Supplementary Table 1.

**Figure 4.**
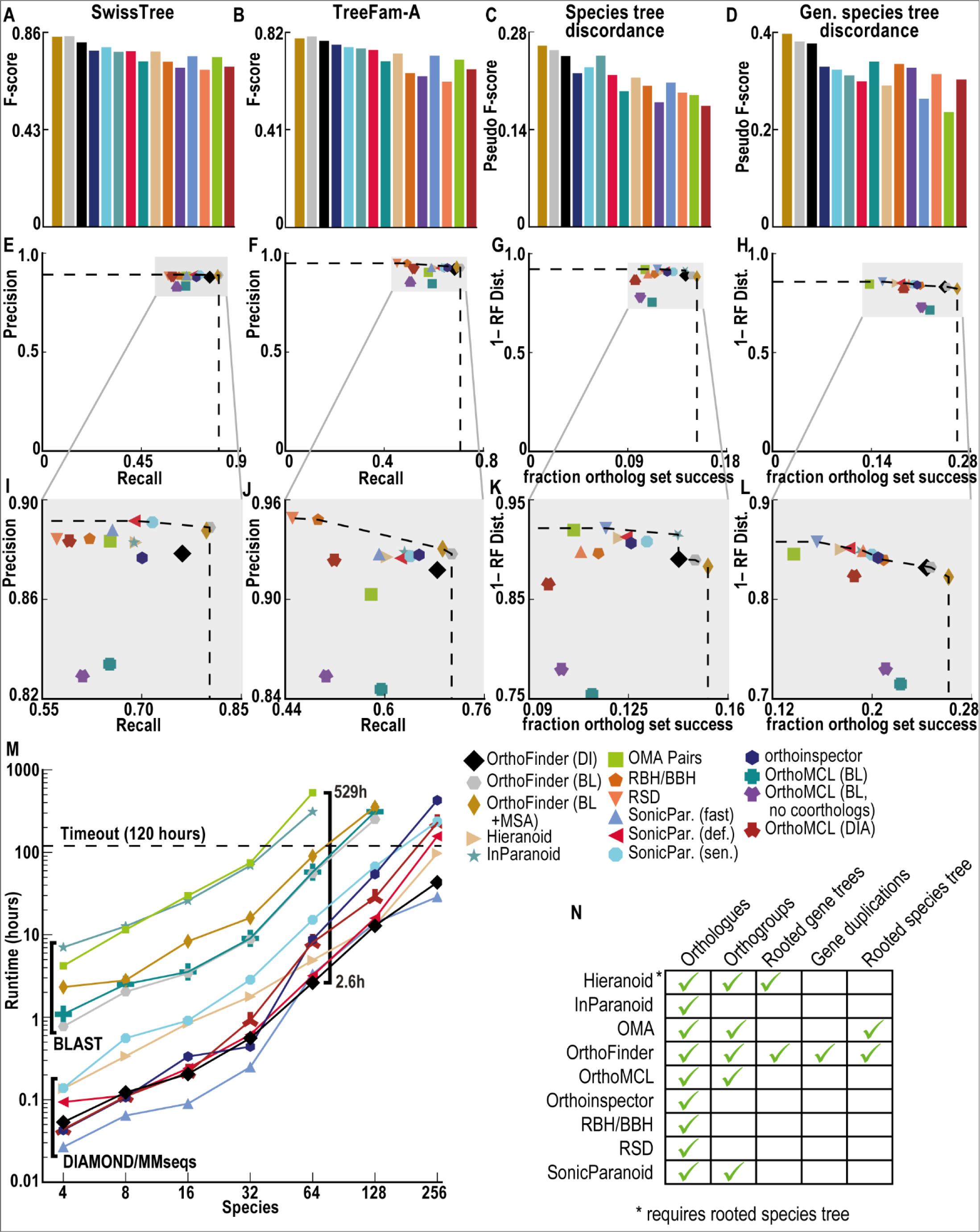
A-L) Quest for Orthologs benchmarks (see [1]) on 66 species across Eukarya, Bacteria and Archaea for ortholog inference methods. Dotted line shows Pareto frontier. Data for graphs are in Supplementary Table 1. A-B) F-score on SwissTree & TreeFam-A tests C-D) ‘Pseudo F-score’ for Species Tree and Generalised Species Tree Discordance Tests. E-F) Agreement of orthologs with SwissTree/FreeFam-A trees G-H) Species tree & generalised species tree discordance tests. X-axis: percentage of randomly selected genes with predicted orthologs in a predefined set of species. Y-axis: (1 – normalised Robinson-Foulds distance) between gene tree for putative orthologs and the known species tree. I-L) Zoom in of plots E-H. See methods for details of Quest for Orthologs benchmarks. M) Runtime for each method with 4-256 input Fungi proteomes. N) Results returned by methods.

The SwissTree and TreeFam-A tests within *Quest for Orthologs* assess the accuracy of ortholog inference against orthologs from gold-standard trees. For these tests, precision, recall and F-score can be calculated. On these tests the default, fastest version of OrthoFinder was 3%-17% (SwissTree, Fig 4A) and 2%-28% (TreeFam-A, Fig 4B) more accurate than any other method. The other versions of OrthoFinder were a further 1-2% more accurate than default OrthoFinder. No method was consistently second best to OrthoFinder.

For the *Quest for Orthologs* Standard and Generalised Species Tree Discordance Tests (STDT & GSTDT) no ground truth orthologs are known, and the methods are assessed on the percentage of trials in which sets of orthologs are identified across a set of species, and the Robinson-Foulds distance between species tree and the gene tree of the putative orthologs. As such, standard precision, recall and F-score measures cannot be calculated. For these tests a ‘pseudo F-score’ was calculated using percentage of recovered ortholog sets in place of recall and 1 - normalised Robinson-Foulds distance in place of precision (equivalently, the proportion of bipartitions in agreement between the species tree and the putative orthologs tree). On both STDT and GSTDT all versions of OrthoFinder had an equal or higher pseudo F-score than all version of all other methods. The default, fastest version of OrthoFinder was 0%-41% (STDT, Fig. 4C) and 11%-60% (GSTDT, Fig 4D) higher scoring than competing methods. The other versions of OrthoFinder were a further 1%-6% higher scoring than the default version. No method was consistently second best to OrthoFinder.

All versions of OrthoFinder, irrespective of algorithmic options, inferred more orthologs (higher recall/recovered ortholog sets) than any other tested method at a similar level of precision (Fig. 4E-L). Across the four tests, the default and fastest version of OrthoFinder (DIAMOND) achieved between 0.2% (Fig. 4G) and 77.8% (Fig. 4D) higher recall/recovered ortholog sets than competing methods. It achieved precision/ortholog species tree agreement between 3.4% lower (Fig. 4C) and 18% (Fig. 4C) higher than competing methods

In addition to testing OrthoFinder against competitor methods than can be run on raw sequence data, OrthoFinder was also compared to static database methods that involve various levels of human curation. All versions of OrthoFinder, irrespective of algorithmic options, had a higher average F-score/pseudo F-score across the four tests than any of the databases (Supplementary Fig. 2). The default version F-score/pseudo F-score was between 2% and 17% higher than the database methods. OrthoFinder (BLAST + MSA) scored between 5% and 20% higher than the database methods (Supplementary Fig. 2). Thus, although OrthoFinder is fully automated and requires no manual curation, it also achieved higher accuracy than curated online database methods.

### OrthoFinder is fast and scales well to hundreds of species

To demonstrate the scalability of the OrthoFinder method, it was run on sets of between 4 and 256 fungal species with 16 parallel processes (Fig. 4M). All other publicly available software tools that have been benchmarked on the *Quest for Orthologs* dataset were similarly tested. The default version of OrthoFinder ran in 192s on the 4 species and 1.8 days on the 256 species datasets. In this time, it inferred orthogroups, all gene trees, the rooted species tree, orthologs and gene duplication events (Fig. 4N). Overall, OrthoFinder was the second quickest method, with the fastest method SonicParanoid taking 1.2 days on the same 256 species set. Both OrthoFinder and SonicParanoid scaled well to the largest datasets, both taking less than half the time of the next best method (4.1 days, Fig. 4M).

There was a large range of runtimes across the complete set of methods. Many methods were unsuited to larger species sets, with 64 species being the largest set on which all methods were runnable within the 120 hour (5 day) cut-off. At this point of comparison, the slowest method took 200 times longer to run than OrthoFinder. It should also be noted that no competitor method also provides gene trees or identifies gene duplication events (Fig 4N). Thus not only is OrthoFinder the most accurate method, and the second fastest method, it also provides the largest quantity of phylogenomic information.

### OrthoFinder efficiently and accurately solves the challenge of inferring a rooted species tree from unaligned protein sequence data

Rooted gene trees are required to enable the use of phylogenetic information for ortholog inference, since correct placement of the root is required for the correct dissection of phylogenetic relationship between genes in the tree [21]. However, the vast majority of tree inference methods infer unrooted trees. Gene trees can be correctly rooted given knowledge of the underlying rooted species tree, and thus OrthoFinder first infers and then roots the species tree for the set of species being analysed. OrthoFinder solves these two challenges (species tree inference and rooting) using two algorithms developed specifically for this purpose.

The species tree is inferred from the set of unrooted orthogroup gene trees using STAG [32] and this species tree is rooted using STRIDE [21]. STAG was developed to leverage the vast amount of phylogenetic information already available in the complete set of orthogroup gene trees inferred by OrthoFinder. It was also developed to be robust to high levels of gene duplication and loss that can hamper methods that rely on sets of single-copy orthologs [32]. It outperformed popular species tree inference methods on benchmark data and scaled well to large datasets [32]. Methods for *ab initio* species tree rooting (i.e. without prior knowledge of a suitable outgroup) have received little attention [21]. STRIDE was similarly developed to leverage the complete set of orthogroup gene trees to efficiently determine the root of the species tree and achieved high accuracy on benchmark data [21]. The ability of OrthoFinder to automatically leverage the raw amino acid sequence data to infer the rooted species tree thus enables outgroup rooting of the complete set of orthogroup gene trees for any input set of species and for all gene trees. This is a critical step for enabling phylogenetic orthology inference from gene sequences.

### OrthoFinder implements a novel duplication-loss-coalescent algorithm to account for gene-tree inaccuracies when identifying gene duplication events and orthologs

Given a set of rooted orthogroup gene trees, the final major challenge in accurately dissecting phylogenetic relationships between genes is to account for incomplete lineage sorting and gene tree error. Existing methods for determining if genes within a gene tree are orthologs or paralogs either had poor accuracy or were unable to scale to the number and size of the orthogroup gene trees that must be analysed. Thus to address this challenge, a novel, scalable algorithm based on the duplication-loss-coalescent model was developed (see methods).

To demonstrate the relative performance characteristics of this method, it was applied to two independent simulated datasets [31, 33] and compared to four popular, comparable methods: Notung [29], GSDI Forester [28], DLCpar (full and search) [31] and the species-overlap method [30]. In terms of accuracy, the novel OrthoFinder method out-performed all methods other than DLCpar (full) (Fig. 5A, Supplementary Table 2). However, DLCpar (full) was unable to analyse realistic sized species datasets. For example, while the OrthoFinder method was able to analyse the complete set of 18651 orthogroup gene trees (948449 genes) from 128 fungal species in 141 seconds, DLCpar (full) was unable to process a considerably smaller, 4 species dataset (2259 trees, 12958 genes) in 120 hours (Fig. 5B). Thus, OrthoFinder is the most accurate method that is scalable to realistic datasets. This algorithm enables accurate interrogation of orthogroup gene trees in a manner that can analyse thousands of gene trees across hundreds of species in minutes on standard computing hardware (Fig. 5B).

**Figure 5.**
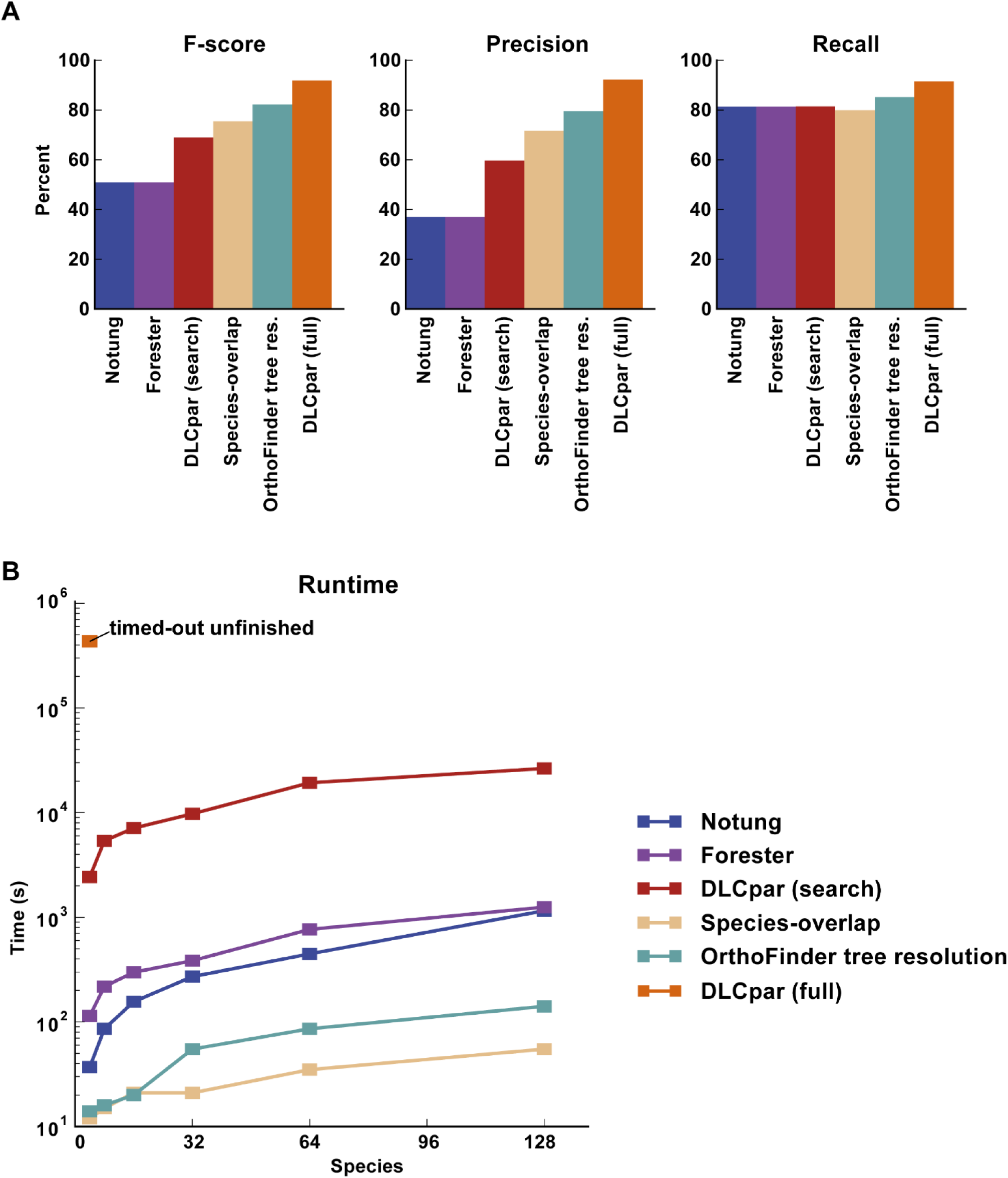
A) Duplication inference accuracy on simulated gene trees. B) Runtime to analyse all trees from the 4 to 128 species Fungi datasets (see methods), a maximum time of 120 hours (4.3×10^5^ seconds) was allowed. DLCpar (full) did not complete the smallest dataset in this time limit and so only the lower bound for the first time point is shown.

## Discussion and Conclusions

Phylogenetic relationships between gene sequences are defined by their relationship in a gene tree in the context of a species tree. Due to the complexity of conducting phylogenetic orthology inference from raw gene sequences, multiple methods have been developed to bypass phylogeny and approximate phylogenetic relationships from heuristics on pairwise sequence similarity scores. Such approximations are subject to common errors that are avoidable by analysis of phylogenetic trees of gene sequences. Here, we present a substantial update to OrthoFinder that provides the first fully phylogenetic orthology inference method.

From testing on community standard benchmarks we demonstrate that OrthoFinder is the most accurate orthology inference method available. Furthermore, we show that by taking a phylogenetic approach OrthoFinder provides substantial additional information (including rooted gene trees, rooted species trees and gene duplication events) that are not provided by heuristic methods. Thus, OrthoFinder is the most accurate and most data-rich orthology inference method for comparative genomics.

The only input required for OrthoFinder is the set of amino acid sequences from protein coding genes for the species of interest. The default parameters for OrthoFinder are optimised for speed and scalability and enable the combined analysis of hundreds of species on commonly available computer resources. However, OrthoFinder is also designed with the expert user in mind and intermediate steps in the algorithm can be substituted with any preferred method for multiple sequence alignment and tree inference should the user wish. We illustrate the time-accuracy trade-off associated with changes in internal steps of the algorithm, and show that the fastest and least accurate implementation OrthoFinder is still more accurate than any other orthology inference method.

## Methods

### OrthoFinder workflow

A gene tree is the canonical representation of the evolutionary relationships between the genes in a gene family. Thus, ortholog inference from gene trees is an important goal. However, no automated software tools are available that provide genome-wide ortholog inference from gene trees. A number of challenges had to be addressed to enable this. These included: the efficient partitioning of genes into small, non-overlapping sets such that all orthologs of a gene are contained in the same set as the original gene; scalable and accurate inference of gene trees from these gene sets; automatic rooting of these gene trees without a user-provided species tree; and robust ortholog inference in the presence of imperfect gene-tree inference. The OrthoFinder workflow was designed to address each of these challenges and is described in detail below.

By default, OrthoFinder infers orthologs from orthogroup trees (a gene tree for the orthogroup) using the steps shown in Figure 1. Input proteomes are provided by the user using one FASTA file per species. Each file contains the amino acid sequences for the proteins in that species. Orthogroups are inferred using the original OrthoFinder algorithm [10]; an unrooted gene tree is inferred for each orthogroup using DendroBLAST [23]; the unrooted species tree is inferred from this set of unrooted orthogroup trees using the STAG algorithm [32]; this STAG species tree is then rooted using the STRIDE algorithm by identifying high-confidence gene duplication events in the complete set of unrooted orthogroup trees [21]; the rooted species tree is used to root the orthogroup trees; orthologs and gene duplication events are inferred from the rooted orthogroup trees by a novel hybrid algorithm that combines the ‘species-overlap’ method [30] and the Duplication-Loss-Coalescent model [31] (described below); and comparative statistics are calculated. All major steps of the algorithm are parallelised to allow optimal use of computational resources. Only orthogroup inference was provided in the original implementation of OrthoFinder [10]; all other subsequent steps are new, and described below.

### Use of orthogroups for gene tree inference

Orthologs are the set of genes in a species-pair descended from a single gene in the last common ancestor of those two species. An orthogroup is the set of genes from multiple species descended from a single gene in the last common ancestor (LCA) of that set of species. Thus an orthogroup is the natural extension of orthology to multiple species.

For ortholog inference, orthogroups are the optimum partitioning of genes for gene tree inference: An orthogroup is the smallest set of genes such that, for all genes it contains, the orthologs of these genes are also in the same set. Since gene tree inference scales super-linearly in the number of genes, partitioning genes into the smallest possible sets is the most efficient way of constructing a set of gene trees that encompass all orthology relationships. Although partitioning genes into larger sets (e.g. gene families containing gene duplication events prior to the LCA) would decrease the number of gene trees to be inferred, the super-linear scaling of gene tree inference would result in a longer overall runtime for the complete set of trees. The original OrthoFinder orthogroup inference method is still the most accurate method on the independent Orthobench test set [10] and thus is used for this step.

### Customisable steps in the OrthoFinder method

There are two customisable steps in the OrthoFinder method. 1) The sequence search method. 2) The orthogroup tree inference method. The default option for step one is DIAMOND [5]. The default option for step 2 is DendroBLAST [23]. The default options are recommended by the authors as they are fast and achieve high accuracy on the Quest for Orthologs benchmarks [1] (Fig. 4A-D). However, the user is free to substitute any alternative methods for these steps. Currently supported methods for step 1 include BLAST [4] and MMseqs2 [6]. Similarly, any combination of multiple sequence alignment and tree inference method can be substituted in for step 2. For illustrative purposes, the default multiple sequence alignment method is MAFFT [34] and the default tree inference method is FastTree [24], this combination is benchmarked above. It is impossible for the authors to test all possible combinations of multiple sequence alignment and tree inference methods, and the selected methods were chosen because of their speed and scalability characteristics [24, 34]. OrthoFinder provides the flexibility for the user to select their preferred method. More accurate multiple sequence alignment and tree inference methods should give more accurate ortholog inference and many studies exist comparing the accuracy and runtime characteristics of the available methods [35, 36]. A user-editable configuration file is provided in JSON format that allows new sequence search, multiple sequence alignment and tree inference methods to be added to OrthoFinder. To facilitate the trialling of alternative multiple sequence alignment and tree inference methods OrthoFinder provides the option to restart an existing analysis after the orthogroup inference stage. This skips the requirement to compute the all-versus-all sequence search and orthogroup inference and thus accelerates testing of different internal steps.

### Species tree inference and rooting

The rooted species tree is required in order to identify the correct out-group in each orthogroup tree, as correct gene tree rooting is critical for the orthology assessment from that tree [21]. Since orthogroups can potentially contain any subset of the species in the analysis, it is not sufficient to simply know the out-group for the complete species set. Instead, the complete rooted species tree is required. If the user knows the rooted species tree for the set of species being analysed then it is recommended to specify this tree manually at the command line to remove the possibility of species tree inference error. Such a tree can be provided as a Newick format text file. In the event that a species tree is not provided (or not known), then OrthoFinder automatically infers it.

Sets of one-to-one orthologs that are present in all species are often used for species tree inference, however in real-world large-scale analyses these can be rare [32]. A new algorithm, STAG (Species Tree from All Genes), was developed to allow species tree inference even for species sets with few or no complete sets of one-to-one orthologs present in all species [32]. Without this algorithm, species tree inference could fail if there were no sets of one-to-one orthologs present in all species. STAG infers the species tree using the most closely related genes within single-copy or multi-copy orthogroups. In benchmark tests, STAG [23] had higher accuracy than other leading methods for species tree inference; including maximum likelihood species tree inference from concatenated alignments of protein sequences, ASTRAL [37] & NJst [38].

The STRIDE algorithm (Specie Tree Root Inference from Duplication Events) [21] is used to root the species tree in OrthoFinder. STRIDE was developed to enable rooting of the species tree using only information available in the set of gene trees. STRIDE does this by identifying the set of well-supported in-group gene duplication events in the complete set of unrooted orthogroup orthogroup trees, and using these events to infer a probability distribution over an unrooted STAG species tree for the location of its root. Similarly to STAG, STRIDE has been shown to correctly identify the correct root of the species tree in multiple large-scale molecular phylogenetic data sets spanning a wide range of timescales and taxonomic groups [21].

### Gene tree rooting

Tree inference methods infer unrooted gene trees. A gene tree must be correctly rooted in order for it to show the correct evolutionary history of the gene family and thus to allow correct ortholog inference. The orthogroup trees could contain any subset of the input species. In general the rooted species tree, inferred as described above, can be used to root the orthogroup trees by identifying the out-group clade in each orthogroup tree and placing the root on the branch separating this out-group from the remaining genes.

However, species tree and gene tree topologies can arise in which this simple approach will not work and so a robust generalisation of this outgroup rooting method is required in order to be able to root any potential gene tree. Firstly, in the species tree the out-group could consist of a single species or multiple species. Secondly, in the gene tree the genes from the out-group could be in a monophyletic clade or there may be no bipartition in the tree that separates all the genes from the out-group from all remaining genes. Thirdly, a gene duplication event could have occurred in the gene tree prior to the divergence of the out-group from the remaining species. Thus, the most ancient bipartition of the gene tree would be a gene duplication event separating the genes into two clades rather than a bipartition separating the out-group from the in-group. Such a gene tree should be rooted on this bipartition. Both of these two descendant clades could then potentially contain genes from both the out-group and in-group species. Thus, there will be no bipartition in such a tree that separates the genes of the out-group species from the genes of the in-group species.

The algorithm used by OrthoFinder searches for the correct bipartition on which to place the root. For each bipartition in the gene tree it calculates two scores. The first, S_AD_, quantifies how well the bipartition corresponds to an ancient duplication prior to the divergence of the species. The second, S_IO_, quantifies how well the bipartition corresponds to the divergence of the out-group species from the in-group species. Both S_IO_ and S_AD_ range between 0 and 1. Let O be the set of species in the out-group and I be the set of species in the in-group. For a bipartition in the unrooted gene tree, let A be the set of species with genes on one side of the bipartition and let B be the set of species with genes on the other side of the bipartition. Then

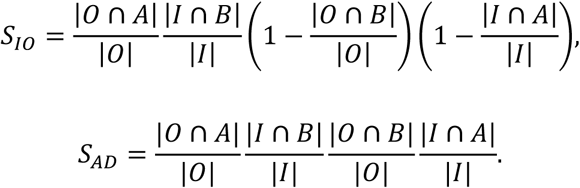

Each of the four terms in these equations quantifies the proportion of in-/out-group species the bipartition correctly includes/excludes from clade A/B of the gene tree (giving the 2^3^=8 terms in total across the two equations). The bipartition with the highest score for either S_IO_ or S_AD_ is the optimal root for the gene tree using this measure.

The effectiveness of these scores at identifying the correct root can be seen by considering the following. A bipartition with a value of 1 for S_IO_ implies that it perfectly divides the tree into an in-group and out-group, and implies a value of 0 for S_AD_ for all bipartitions in the tree (thus there are no potential bipartitions corresponding to an ancient duplication). This is the correct bipartition on which to root the tree since it separates the in-group from the out-group genes. Conversely, a bipartition with a value of 1 for S_AD_ implies that the bipartition is a duplication event before the divergence of any of the species, with all species present for both duplicates. It implies a value of 0 for S_IO_ for all bipartitions in the tree (thus there is no bipartition that corresponds to a first speciation event that splits the genes into an out-group clade and an in-group clade). The highest value for either S_IO_ or S_AD_ across the tree shows that the corresponding bipartition is close to one of these perfect cases and is the best root for the gene tree.

### Ortholog inference and identification of gene duplication events from gene trees

A number of methods were considered for distinguishing orthologs from paralogs in gene trees. Duplication and loss reconciliation (e.g. Forester [28], Notung [29]) uses a rooted species tree and rooted gene tree to determine if each node in the gene tree is a speciation or a duplication event. Genes that diverged at a speciation event are orthologs whereas those that diverged at a duplication event are paralogs. DLCpar [31] uses a model for duplication-loss-(deep) coalescent (DLC) that addresses incongruence between the gene and species trees to increase accuracy. It exists in two versions which we label DLCpar (full) and DLCpar (search). DLCpar (full) considers the complete space of possible reconciliations to find the maximum parsimony solution under the DLC model, but can have large run times even for relatively small gene trees. DLCpar (search) instead employs an iterative search for a locally optimal solution, which can differ from the globally optimal solution. A third approach, here referred to as the species-overlap method is employed in a number of ortholog databases [18, 30] and was originally described in a method for determining orthologs of human genes [30]. In this method, nodes in the gene tree are identified as duplication nodes if the sets of species below its child nodes overlap, otherwise the node is a speciation node. Genes that diverged at a speciation node are orthologs, those that diverged at a duplication node are paralogs.

These methods were tested on the fungal orthogroups (in parallel, using 16 cores) to determine their runtime on sets of typical orthogroup trees derived from sets of between 4 and 128 species. Our implementation of the species-overlap method was the fastest, taking 55s to analyse the largest datatset (Fig. 5). This dataset consisted of the 18651 orthogroup trees containing 948449 genes and corresponded to the complete set of orthogroup trees for the 128 fungal species. Notung and Forester were both 21-23 times slower, DLCpar (search) was over 500 times slower. DLCpar (full) was unable to complete the analysis of the smallest input dataset in 120 hours and so was not tested on any of the larger datasets. To put this time in context, all steps in the OrthoFinder algorithm for this dataset collectively take less than 4 minutes in total (i.e. orthogroup inference, gene tree inference, species tree inference, species tree rooting, gene tree rooting).

To compare the accuracy of the above methods, they were each tested for their precision and recall in identifying gene duplication events on simulated ‘flies’ and ‘primates’ datasets [31] and a simulated ‘metazoa’ dataset [33]. Since for all methods tested a node in a gene tree is either a duplication or speciation event, the identification of all gene duplication events is equivalent (by complementation) to the identification of all speciation events. Thus, overall accuracy at identifying gene duplication events is equivalent to the overall accuracy at identifying orthologs. The most accurate method on the simulated data was DLCpar (full) with and F-score of 91.8% followed by the species-overlap method with an F-score of 75.5%.

Since DLCpar (full) was the most accurate method on the simulated datasets but was unsuitable for analysing gene trees with more than four species a novel hybrid algorithm was developed. This aimed to combine the strengths of the highest accuracy DLCpar (full) method with simplifications from the species-overlap method to achieve high accuracy in a reasonable run time.

In the DLC model, clades of genes containing no duplicates are analysed to find the most parsimonious reconciliation with the species tree. This is required since the goal for DLCpar is a complete reconciliation of the gene tree with the species tree. However, in species-overlap method, clades of single-copy genes are identified as orthologs without further analysis of the topology of their relationship. This assumption is reasonable, since trees of single-copy orthologs are frequently topologically distinct from the species tree. For example, in an analysis of 1030 gene trees of one-to-one orthologs from 23 fungi species all 1030 gene trees were topologically distinct from each other and from the species tree [39]. Analysis of such clades under the DLC model is likely to be computationally costly with no benefit in terms of accuracy of ortholog inference.

On the other hand, when a gene duplication event has occurred it is important to accurately identify the genes affected by this event since the location of the event determines which genes are orthologs and which are paralogs. In the hybrid algorithm developed for OrthoFinder these nodes, for which there is evidence of a gene duplication event through overlapping species sets, are analysed under the DLC model. The DLC model is used to attempt to find the most parsimonious interpretation of this node in terms of which genes diverged at the gene duplication event and which diverged as a speciation event.

As described, this method would still require exploring a large search space for the nodes under consideration and the reduction in runtime would not be significant. Thus, to accelerate the process duplication and loss events are inferred directly using the species-overlap method. A duplication event is inferred from an overlap in the species sets below a node and a loss event is inferred by the presence of a gene from a species in one of the descendant clades but not in the other. The analysis can then be accelerated by classifying a node according the species-overlaps of its subclades up to a maximum total topological depth of two below the node being analysed (clades O., Supplementary Fig. 3A). The possible sub-cases for the overlaps between these clades have been enumerated (Supplementary Fig. 3B). For each sub-case, the most parsimonious interpretation under the DLC model has been pre-calculated (Supplementary Fig. 3C) and can thus be corrected without need for a topology search.

The algorithm implemented in OrthoFinder is as follows. A post-order traversal of the orthogroup tree is performed (a node is not visited until all its descendant nodes have been visited), analysing each node of the orthogroup tree in turn. A given node is analysed to identify if the species sets below its child nodes overlap. If there is an overlap, the smallest sub-clade below each child nodes that contains the complete set of overlapping species is identified up to a maximum total topological depth of two below the node (clades O., Supplementary Fig. 3A). The node is assigned to the corresponding sub-case (Supplementary Fig. 3B). If a more parsimonious interpretation of the sub-case is available under the DLC model then the sub-tree below the node is rearranged to match this interpretation (Supplementary Fig. 3C). After the node has been analysed, the next node in the post-order traversal is analysed. Note, the choice of a post-order traversal allows the traversal to be continued unimpeded despite any such rearrangements below the node being analysed. The resulting gene trees are referred to as ‘resolved’ gene trees and correspond to the ‘locus tree’ under the DLCpar model [31]. Orthologs and gene duplication events are determined from the resolved gene tree according to the species overlap method.

Although only a single traversal of the tree is employed, rather than the iterative search and rearrangement employed by DLCpar, the post-order traversal enables more parsimonious interpretations of child clades below a node to be identified prior to the analysis of the parent node. Thus, analysis of sub-trees below a node inform the subsequent analysis of the node itself. In theory, nodes could be categorised to sub-cases based on overlaps of clades at a greater topological depth than that employed here. This conservative approach was taken since the number of subcases increases exponentially and a total topological depth of two proved sufficient to achieve a higher accuracy for the method compared to the simple species-overlap. The analysis of clades to this depth proved sufficient to increase the F-score from 72% with just the species-overlap method to 80% with the hybrid algorithm (Fig. 5A). The pre-calculated solutions for each sub-case removed the need for costly, iterative search using random (i.e. unguided) tree rearrangement operations thus accelerating the analysis considerably. The hybrid algorithm was able to analyse the complete set of orthogroup trees for the 128 fungi species in 141s, this was 8 times faster than Notung and 187 times faster than DLCpar (search) (Fig. 5B). The hybrid method also outperformed both methods in terms of accuracy (Fig. 5B). Note that the species tree is not required for the hybrid model used by OrthoFinder. The only use of the species tree is in determining the root for each orthogroup tree.

### Simulation Tests of OrthoFinder Gene Duplication Event Inference Accuracy

The tests for gene duplication event inference accuracy were performed on the simulated ‘flies’ and ‘primates’ dataset from [31] and a simulated ‘metazoa’ dataset from [33]. To model real data, the flies and primate datasets used known species trees, parameters for divergence times, duplication rates, loss rates, population sizes, and generation times. Trees were simulated with varying effective population sizes and duplication rates so as to model incomplete lineage sorting [31] [33]. The flies dataset consisted of 12,000 trees with 12 species and 12,032 gene duplication events. The primates dataset consisted of 7,500 trees with 17 species and 16,066 gene duplication events. The metazoa dataset intended to emulate the complexity of real data by using heterogeneity in rates of duplication and loss, a complex model of sequence evolution and then inferring trees with a homogenous, simple model [33]. It consisted of 2,000 gene trees with 40 species and 4,967 gene duplication events. For comparison, Forester [28], Notung [29], DLCpar (full), DLCpar (search) [31] and the overlap algorithm (i.e. without OrthoFinder’s tree resolution) were also tested.

All methods were provided with the input rooted gene tree and, where appropriate, the rooted species tree (Forester, Notung, DLCpar). The duplication cost used by Notung was set to 1.0. No other parameters required specification for any of the other methods. The rooted gene trees were provided as part of the simulated data for the flies and primates datasets. Multiple sequence alignment (MSA) files were provided for the metazoa dataset. For this dataset gene trees were inferred from the MSAs using FastTree so as to also include a potential level of tree inference error and were rooted with reconroot [31]. The OrthoFinder rooting algorithm was not used so as to avoid inadvertently biasing the results in favour OrthoFinder. All methods were provided with the same input rooted gene trees. The complete set of gene duplication events identified by each of the methods were compared against the ground truth gene duplication events. An inferred gene duplication was identified as correct if the two sets of genes observed post-duplication exactly matched the two sets of genes post-duplication from the ground truth data.

The performance testing of the methods for identifying gene duplication events was performed on the orthogroup trees from the 4-to 128-species fungi datasets as inferred by OrthoFinder with default parameters. The commands for Forester, Notung and DLCpar were run in parallel using GNU Parallel [40] using 16 threads on these gene trees. The OrthoFinder method was run via the “scripts/resolve.py” program included as part of the OrthoFinder distribution. To allow testing, the species-overlap method was also implemented in OrthoFinder and was run using the same program with the option “--no_resolve”.

### Ortholog Benchmarking

Orthogroup inference accuracy of OrthoFinder has already been tested using the independent Orthobench dataset [11]. This showed it to be the most accurate method tested [10]. The community developed ‘Quest for Orthologs’ benchmarks [1] were used to assess the accuracy of the newly developed OrthoFinder ortholog inference. OrthoFinder was tested using the default method (DIAMOND sequence search and DendroBLAST trees, no additional options). It was also tested with the BLAST replacing DIAMOND (options: “-S blast”), and with both BLAST search and multiple sequence alignment and maximum likelihood tree inference (options: “-S blast -M msa”). In the latter MAFFT [34] and FastTree [24] were used for multiple sequence alignment and tree inference as described above. For each of these three cases, OrthoFinder was run on the 64 reference proteomes of the Quest for Orthologs test set with a single command (“-f Proteomes/” + options) and the inferred orthologs were submitted to the Quest for Orthologs webserver for benchmarking.

The Quest for Orthologs benchmarks are described in detail in [1]. The species tree discordance test (and the generalised version of this test) both consider a set of species partitioned into clades with a known species tree topology connecting the clades. The benchmarking consists of a repeated test. For one of the clades of species a gene is selected at random for each instance of the test. If the orthology inference method under scrutiny predicts an ortholog for that gene for at least one species from each of the remaining clades then the test is recorded as a ‘successful ortholog set’. For each successful ortholog set an MSA is constructed and a gene tree inferred using RAxML [27]. The normalised Robinson-Foulds (RF) distance is calculated between this tree and the known species tree. The result of the benchmark is the percentage of successful ortholog sets and the average RF distance for the successful sets. A higher percentage success and a lower average RF distance indicates a better ortholog inference method under this test. The SwissTree [41] and TreeFam-A [42] tests compare the ortholog precision and recall compared with the orthologs inferred from a set of manually curated SwissTree and TreeFam-A gene trees. Note that OrthoFinder can be penalized by an increased RF distance in these tests in cases where OrthoFinder judges it more parsimonious that genes are orthologs under the duplication-loss-coalescent model even if the gene tree conflicts with the species tree. The benchmarks treat all such differences as an error, rather than considering what is the most parsimonious explanation for the conflict. This is an important point, as it is expected that gene trees will contain tree error (else species trees could be accurately inferred from a gene tree of any single copy gene). In general, genes trees do not match the species tree and in an exemplar analysis of 1030 gene trees of one-to-one orthologs from 23 fungi species all 1030 gene trees were topologically distinct from each other and from the species tree [39]. Thus, this benchmarking method likely penalises all methods to some extent for making correct ortholog predictions.

The full set of benchmarks, the input files and the ortholog inference results can be seen online at http://orthology.benchmarkservice.org/ and are shown in Fig. 4A-L of the main paper. The complete datasets are available to download from Zenodo research archive at https://doi.org/10.5281/zenodo.1481147.

### Performance Testing

We constructed sets of fungal proteomes of increasing size for performance testing. Ensembl Genomes was interrogated on 6^th^ November 2017 using its REST API [43] to identify all available fungal genomes. To achieve an even sampling of species we selected one species per genera and excluded genomes from candidate phyla or phyla with fewer than 3 sequenced genomes. This gave a set of 272 species which were downloaded from the Ensembl FTP site [44]. We created datasets of increasing size by randomly selecting 4, 8, 16, 32, 64, 128 and 256 species such that the last common ancestor was the same for each dataset. Each dataset was analysed using a single Intel E5-2640v3 Haswell node (16 cores) on the Oxford University ARCUS-B server using 16 parallel threads for OrthoFinder with DIAMOND (arguments: “-S diamond -t 16 -a 16”). The complete datasets for all analysed species subsets are available for download from Zenodo at https://doi.org/10.5281/zenodo.1481147. All methods submitted to Quest for Orthologs that provided a user-runnable implementation of the method were tested on the same Fungi datasets and same ARCUS-B server nodes and run in parallel using 16 threads (when supported by the method).

### Chordata Dataset

The data for the OrthoFinder analysis of the ten chordata species for the illustration of the results of an OrthoFinder analysis (Fig. 2A-H) are provided in the Zenodo archive https://doi.org/10.5281/zenodo.1481147. This includes the input proteomes, the OrthoFinder results and the script used to generate the figures from the results. OrthoFinder was run with default settings (DIAMOND sequence search and DendroBLAST gene trees).

## Declarations

### Ethics approval and consent to participate

Not applicable

### Consent for publication

Not applicable

### Availability of data and material

The OrthoFinder source code and executables are available at https://github.com/davidemms/OrthoFinder. A compressed archive of all data is available at the

Zenodo research data archive at https://doi.org/10.5281/zenodo.1481147.

### Competing interests

The authors declare that they have no competing interests.

### Funding

This work was supported by the European Union’s Horizon 2020 research and innovation program under grant agreement number 637765. SK is a Royal Society University Research Fellow.

### Authors’ contributions

DE and SK conceived and designed the project. DE developed the algorithms. DE and SK discussed the results and wrote the manuscript. All authors read and approved the final manuscript.

## Acknowledgements

The authors would like to acknowledge the use of the University of Oxford Advanced Research Computing (ARC) facility in carrying out this work.

## Supplementary Figure Legends

**Supplementary Figure 1.**
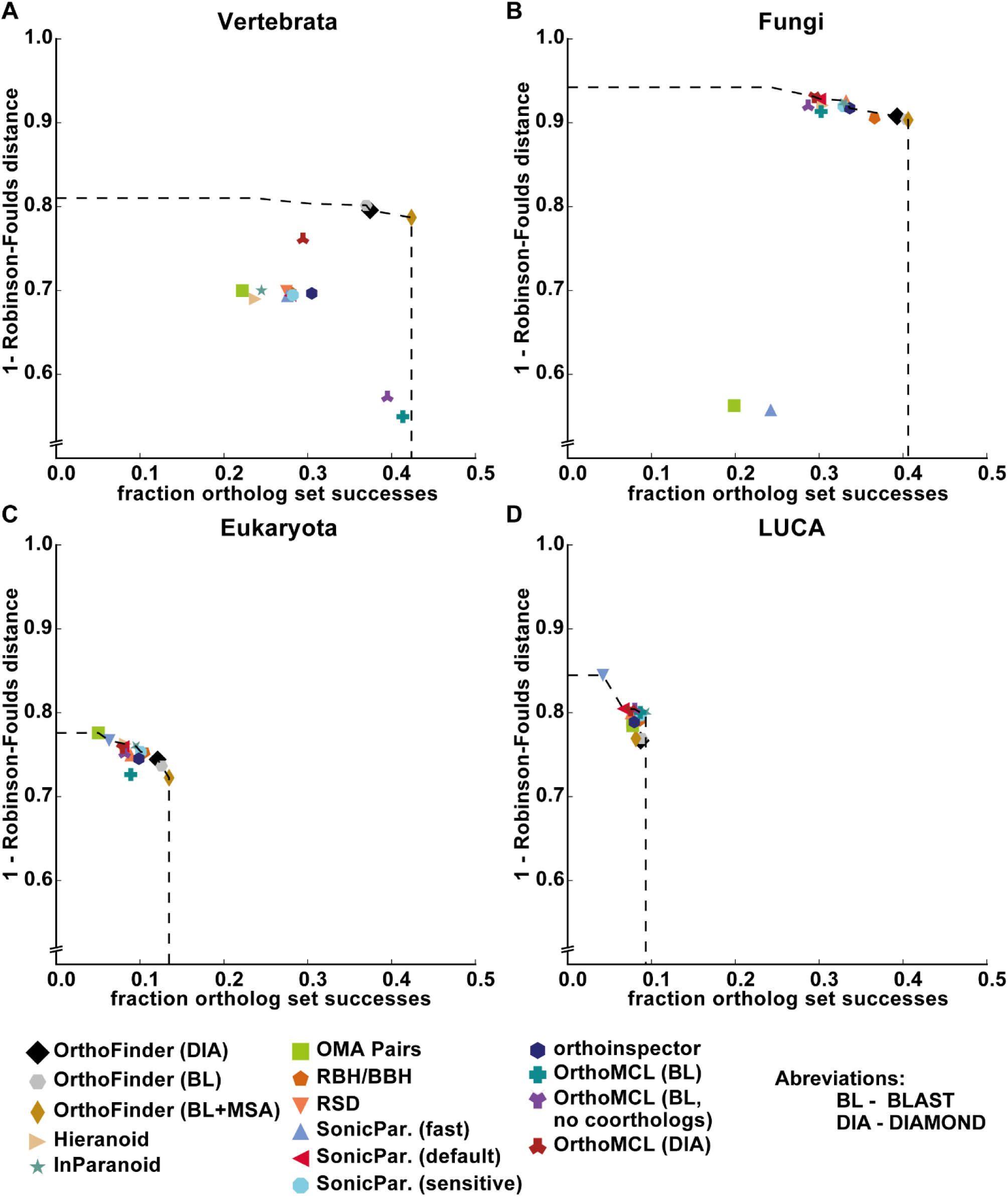
Results of the Generalised Species Tree Discordance tests from the Quest for Orthologs benchmarks for four clades of species. A) Vertebrata B) Fungi C) Eukaryota D) LUCA (Last Universal Common Ancestor, a species set covering bacteria, archaea and eukaryotes). See methods for description of Quest for Orthologs benchmarks.

**Supplementary Figure 2.**
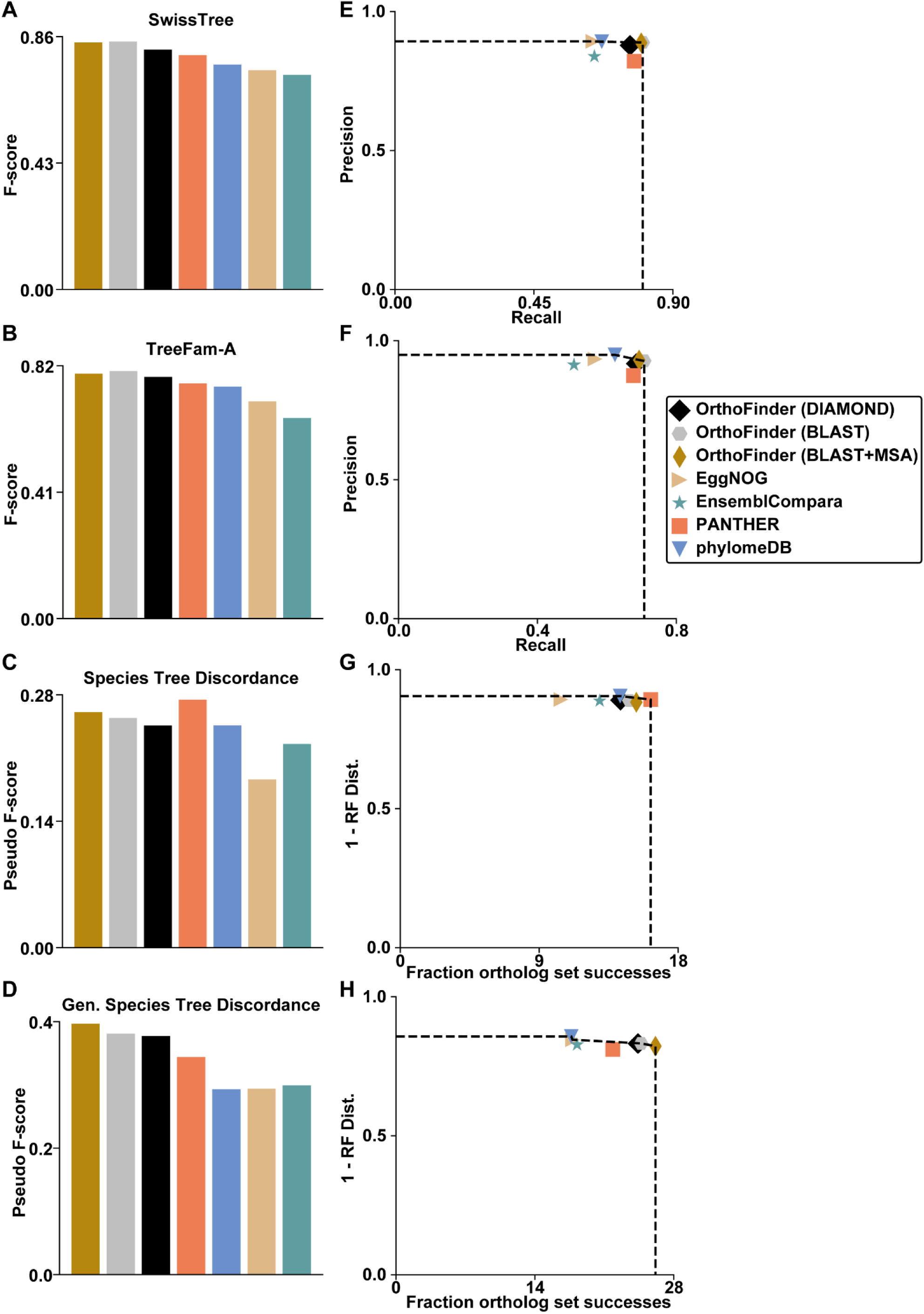
Quest for Orthologs benchmark results for OrthoFinder versus online databases. A-B) F-score versus orthologs from gold standard trees C-D) Pseudo F-score on species tree discordance test E-F) Precision & Recall versus gold standard trees G-H) 1 minus normalised Robinson-Foulds (RF) distance & fraction of ortholog set successes on species tree discordance tests.

**Supplementary Figure 3.**
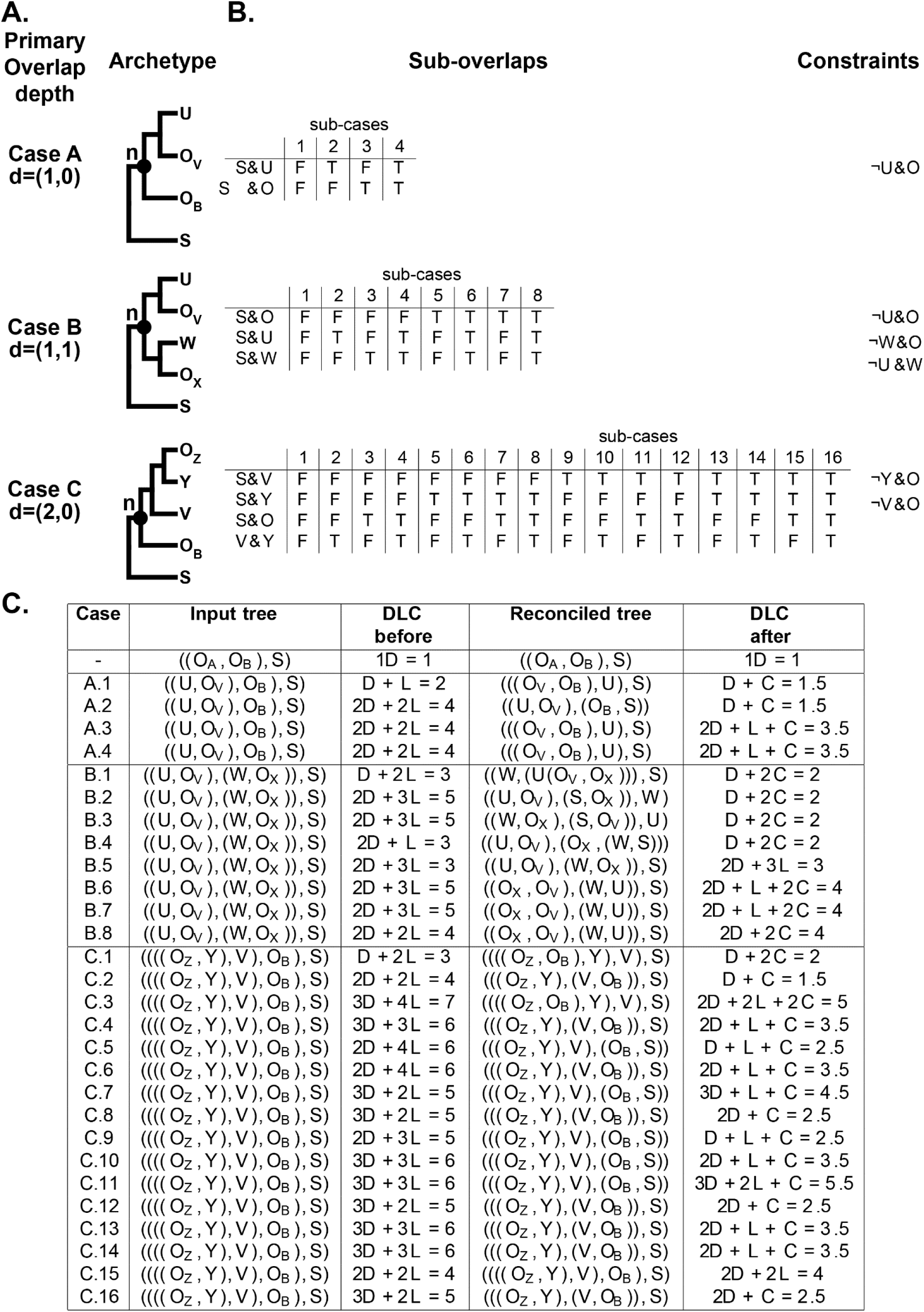
Deterministic tree reconciliation applied to nodes within a gene tree. If the sets of species below a node, n, overlap the node is analysed to find if there is a more parsimonious interpretation of the subtree under the duplication, loss, deep-coalescence (DLC) model. The analysis is done in the context of the sister clade, S, and the descendant clades, U, V, W, X, Y, Z and O. These clades may contain single or multiple genes. The overlapping clades under the two descendants of n are identified, down to a total combined depth of 2 (so that the problem remains tractable). Each case has a number of possible sub-cases according to whether the species sets in the clades overlap. The notation X&Y means that the species set for the genes in clade X overlap with the species sets in the clade Y. In order for a node to fit an archetype (the overlap in n’s descendants is in the clades ‘O’), constraints arise on some of the sub-overlaps. T=True, F=False.

